## Supplemental files for "Regulation of Myoepithelial Differentiation in 3-Dimensional Culture"

**Table S1: Antibody Dilutions & Applications**

| Antibody | Company | Catalog # | Dilution | Application |
|---|---|---|---|---|
| **Primary Antibodies** | | | | |
| αSMA | Sigma Aldrich | A5228 | 1:1000 | ICC |
| TAZ | Sigma Aldrich | T4077 | 1:200 | ICC |
| Calponin | Abcam | Ab46794 | 1:600 | ICC |
| **Secondary Antibodies** | | | | |
| AF568 Donkey anti-Rabbit | Alexa-Fluor | A10042 | 1:1000 (2D)<br>1:500 (3D) | ICC |
| AF488 Donkey anti-Mouse | Alexa-Fluor | A21202 | 1:1000 (2D)<br>1:500 (3D) | ICC |
| DRAQ5 | Cell Signaling Technology | 4084 | 1:1000 (2D)<br>1:500 (3D) | ICC |

**Table S2: qPCR Primer Sequences**

| Gene Product | Strand | Sequence (5'-3') |
|---|---|---|
| αSMA | Forward | GTCCCAGACATCAGGGAGTAA |
| | Reverse | TCGGATACTTCAGCGTCAGGA |
| AQP5 | Forward | AGAAGGAGGTGTGTTCAGTTGC |
| | Reverse | GCCAGAGTAATGGCCGGAT |
| Calponin | Forward | GCACATTTTAACCGAGGTCCT |
| | Reverse | CTGATGGTCGTATTTCTGGGC |
| CTGF | Forward | CTCCACCCGAGTTACCAATG |
| | Reverse | TGGCGATTTTAGGTGTCC |
| CYR61 | Forward | ACCAATGACAACCCAGAGTG |
| | Reverse | AAGTAAATCTGACTGGTTCTGGG |
| Integrin β4 subunit | Forward | ACTCCATGTCTGACGATCTGG |
| | Reverse | GGGACGCTGACTTTGTCCAC |
| TAZ | Forward | GTGTGCCCAATGCACTGA |
| | Reverse | TGACGCATCCTAATCCTCTCTC |
| α-actin | Forward | GGCTGTATTCCCCTCCATCG |
| | Reverse | CCAGTTGGTAACAATGCCATGT |

**Table S3: siRNA sequences**

| Gene Name | Catalog Number | Company | Sequence |
|---|---|---|---|
| WWTR1 (TAZ), Sequence 1 | Custom | Dharmacon | CAGAAUGACUUUAGAGAAUUU |
| WWTR1 (TAZ), Sequence 2 | J-041057-09- 0002 | Dharmacon | CAAUUUAUGUCCACGUUAA |

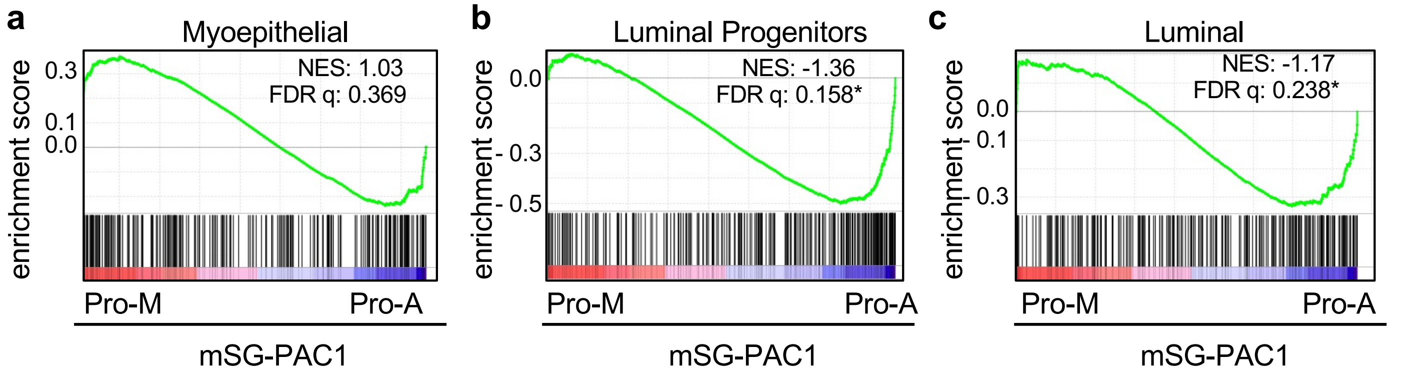


**Supplementary Figure 1. Distinct differentiation programs are engaged in mSG-PAC1 spheroids cultured in Pro-A and Pro-M.** GSEA was performed using gene sets derived from single cell RNA sequencing (scRNAseq) from murine mammary gland for myoepithelial cells and luminal cells from pregnant females and luminal progenitor cells from virgin females. Data sets were downloaded from Supplemental Table 5 data for each cell type[38]. The analysis was performed using the 250 genes with the lowest adjusted p-value. **(a)** Myoepithelial specific genes are modestly, but not significantly enriched in mSG-PAC1 spheroids cultured in Matrigel in Pro-M medium (NES = 1.03, FDR q-value = 0.369). **(b)** Genes specific to luminal progenitor (NES = -1.36, FDR q-value = 0.158) and **(c)** mature luminal cells (NES= -1.17, FDR q-value = 0.238) are significantly enriched mSG-PAC1 spheroids cultured in Matrigel in Pro-A medium (FDR q-value < 0.25 represent significant enrichment.

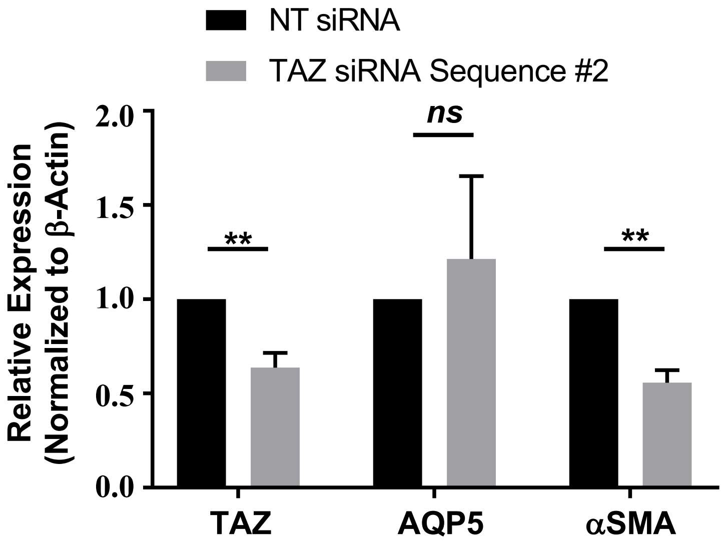


**Supplementary Figure 2.** TAZ depletion using a second siRNA sequence also inhibits the expression of αSMA without significantly affecting the expression of AQP5.

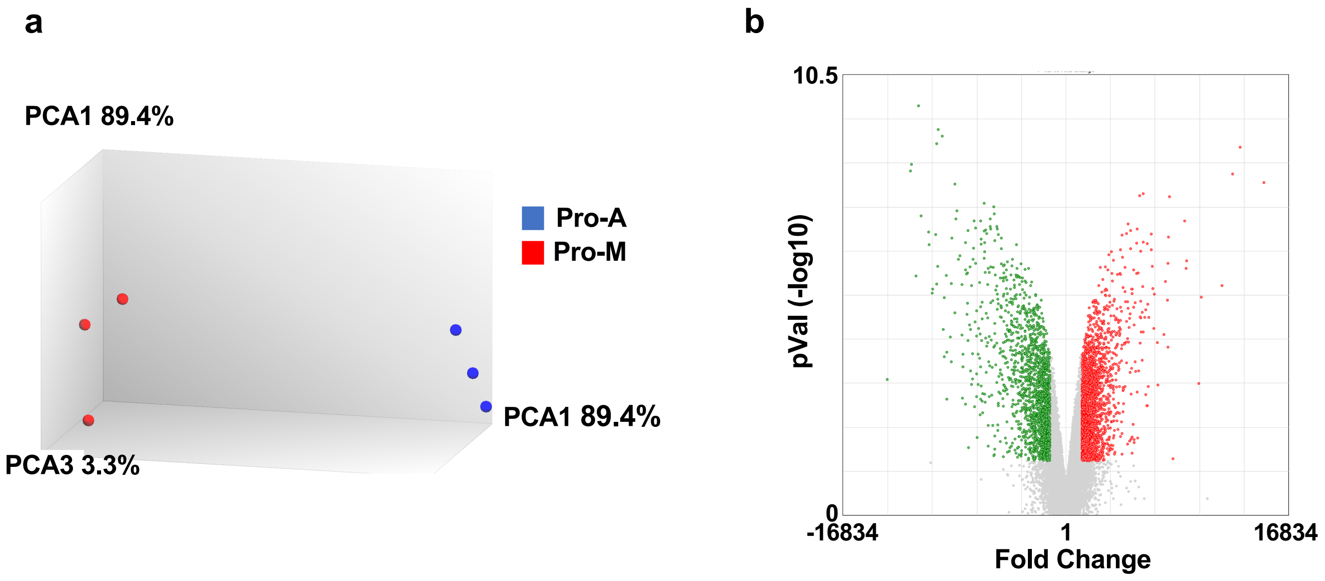


**Supplementary Figure 3. (a)** Principal component analysis (PCA) was performed to examine the variance between samples. **(b)** Volcano plot for differentially expressed genes in spheroids cultured in Pro-M compared to Pro-A medium. Genes increased by a factor of greater than 2-fold are shown in red and those that decreased by a factor of greater than 2-fold are shown in green.
